# A short review of machine learning methods for classifying the outcome of Gestational Diabetes

**DOI:** 10.1101/643833

**Authors:** Shreeya Banerji

**Affiliations:** Department of Statistics, University of Allahabad, Prayagraj, Uttar Pradesh, India

**Keywords:** Diabetes, Data Mining, Classification, Pima Indians

## Abstract

Diabetes mellitus is a growing problem, especially in developing countries. People suffering from diabetes have an increased risk of developing a number of serious health problems. Consistently high blood glucose levels can lead to serious diseases affecting the heart and blood vessels, eyes, kidney, etc. In addition, people with diabetes also have a higher risk of developing infections.

This paper aims to use suitable data mining and classification techniques which include the Logit model, the Probit model, the Classification tree technique, Artificial Neural Networks, Support Vector Machines, Ridge Regression technique and the Least Absolute Shrinkage and Selection Operator(LASSO) in order to determine the best method which can be used to classify the patients as suffering from gestational diabetes or not. The misclassification rate is calculated for different methods and the method having the least misclassification rate is said to be the most suitable to be applied to the given data, which is the PIMA Indians diabetes dataset.

## 1. INTRODUCTION

Diabetes mellitus is a growing problem, especially in developing countries. People suffering from diabetes have an increased risk of developing a number of serious health problems. Consistently high blood glucose levels can lead to serious diseases affecting the heart and blood vessels, eyes, kidney, etc. In addition, people with diabetes also have a higher risk of developing infections. Diabetes can be classified into two main types, Type 1 diabetes and Type 2 diabetes.

Under Type 1 diabetes, the body does not produce enough insulin, and blood glucose levels remain high unless a person takes steps to manage the high blood sugar. Under Type 2 diabetes, which is more common, the body does not respond well to insulin. People with type 2 diabetes have insulin resistance. The body still produces insulin, but it’s unable to use it effectively.

Because of the above mentioned risks, it is very important to develop a predictive model using the risk factors for the development of diabetes. In this paper, the techniques used include the classification tree, artificial neural networks, the logit and the probit models, support vector machines, ridge regression and the least absolute shrinkage and selection operator (LASSO). The R software has been used which is an open source software under the GNU General Public License.

Machine learning is mainly helpful in situations where a given dataset needs to be analysed and conclusions drawn regarding its specific properties, effects of the various variables on the data, etc. Machine learning tools make the analysis of data easier, that is, conclusions can be easily derived which may not be visible directly. A logit model is a type of model that is useful for binary classification when the output is discrete. The logistic function helps to convert the output obtained from the linear combination of the independent variables to lie in between 0 and 1. A probit model works in a similar manner except that is uses the cumulative distribution function (CDF) of a standard normal distribution instead of the logistic function. For probit model as well, the output is obtained in the form of 0 and 1.

The classification tree technique, as the name suggests, is primarily used for classifying datasets in a suitable manner according to the importance of the independent variables and the output is obtained in the form of the predicted classes. The artificial neural network is a method in which the working of the brain is implemented through the use of artificial neurons. It helps in classification by fitting a suitable model using the most appropriate learning rule, with a suitable activation function, weights, and bias, with a given number of hidden layers and nodes which are commonly determined through trial and error.

The support vector machine is a method which uses a suitable algorithm and a separating hyperplane in order to classify the given data. When a hyperplane has a large distance from a point of data in the training set, it may be assumed to be a suitable hyperplane. Ridge regression is a method for fitting a model on data which may be affected by multicollinearity. It uses a penalized loss function having an *l*_2_ penalty, where the sum of the squares of the coefficients of the independent variables are minimized. The Least Absolute Shrinkage and Selection Operator, known as LASSO, also works on data having multicollinearity, but the penalty applied on the loss function is known as the *l*_1_ penalty, where the absolute values of the coefficients of the independent variables are minimized.

## 2. RELATED WORKS

There are several methods which have been used in the study of the classification of the given diabetes dataset. In the medical sector, the classification systems have been mainly used to study the patient’s data and make the predictive models or build a set of rules. Bozkurt, Yurtay, et al. [1] have used different techniques such as the Gini algorithm from decision trees, Artificial Neural Networks (ANN), Distributed time delay networks(DTDNs),Probabilistic neural networks(PNNs) etc. and compared the performance of these classifications. The main objective of the research work by Dilip Kumar Chaubey, Sanchita Paul [2] and others is to provide a better classification of diabetes. Hina S, Shaikh A, Sattar SA.[3] deals with the effects of non-communicable diseases such as stroke, heart disease, cancer, chronic lung cancer and diabetes. They have used data mining techniques which help to obtain information about the data of diabetic patients. Iyer A, Jeyalatha S, Sumbaly. R.[4] use data mining techniques such as decision trees and Naïve Bayes estimator in order to predict the onset of diabetes. The General Regression Neural Network (GRNN) has been used to study the Pima Indians diabetes data in the study by Kayaer, K., and Yildirim, T. [5]. In the paper by Kim Y, et al.[6], the LASSO technique has been applied on the PIMA Indians diabetes dataset by defining a gradient descent algorithm and checking for its convergence to the optimum under the defined regularity conditions.

A study by Koklu et al. [7] deals with the classification of type 2 diabetes using the Weka software package. The Bayesian Network classifier was used to classify patients as being diabetics or not in the paper by Kumari M. et al. [8]. Mohan, V., Usha, S., & Uma, R. [9] have addressed the problems associated with screening for diabetes in developing countries such as India. Various data mining algorithms have been compared by M. Nabi, Pardeek Kumar and others. [10]. They have inspected the execution of four machine learning algorithms namely Naïve Bayes, Logistic Regression, J48 and Random forest to predict the population who are most likely to develop diabetes on Pima Indian diabetes data. The sensitivity of Artificial Neural Networks has been checked in the paper by Olden JD, Jackson DA. [11]. J.W. Smith et al., [12] have used the ADAP learning algorithm to predict the onset of diabetes and compared the results with those of logistic regression and linear perceptron methods. The paper by Stathakis [13] aims to predict the accuracy of different methods for obtaining the number of nodes and hidden layers for the artificial neural networks. The paper by Wu H, Yang S, Huang Z, He J, Wang X. [14] has addressed the problem of improving the accuracy of the prediction model, and to make the model adaptive to more than one dataset.

## 3. METHODOLOGY

### 3.1 Dataset and its description

The PIMA Indians diabetes dataset has been used in the study. All patients in the dataset are females of age 21 years or older, belonging to the Pima Indian tribe near Phoenix, Arizona, USA.

#### Attribute Information for the variables

1. Number of times pregnant- the number of times the patient was pregnant.
2. Plasma glucose concentration-Plasma Glucose concentration is a measure of how much sugar/glucose a person has circulating in their blood.
3. Diastolic blood pressure (mm Hg)- The diastolic pressure is specifically the minimum arterial pressure during relaxation and dilatation of the ventricles of the heart when the ventricles fill with blood.
4. Triceps skin fold thickness (mm)- A measurement used to evaluate nutritional status by estimating the amount of subcutaneous fat.
5. 2-Hour serum insulin (mu U/ml) – Insulin hormone level in the blood, measured in (mu U /ml) where U is defined as the unit of enzyme per milliunit(ml)
6. Body mass index (*weight in kg/(height in m)^2*) -An approximate measure of whether someone is over- or underweight, calculated by dividing their weight in kilograms by the square of their height in metres.
7. Diabetes pedigree function- It provides some data on diabetes mellitus history in relatives and the genetic relationship of those relatives to the patient.
8. Age (years) – Age of the patients.
9. Class variable (0 or 1)- Classifies whether the patient has diabetes (1) or does not have diabetes (0).

### 3.2 Methods used

#### 3.2.1 Logit models

The logistic model or logit model uses a logistic function and models a dependent variable with a binary output. The dependent variable has two possible outcomes in the form of yes/no, win/lose, etc., which are represented by an indicator variable. For the logit model, the log odds ratio for the value labelled as “1” or “success”, which shows the occurrence of the event under study is a linear combination of one or more predictor variables which may or may not be continuous.

#### 3.2.2 Probit models

A probit model is a type of regression model where the dependent variable can only take two values, that is, it is a binary response variable. It works in a manner similar to the logit model, the difference being that of the link function. The probit model uses a probit link function whereas the logit model uses a logit link function. The probit model is useful when dealing with ordinal data.

#### 3.2.3 Classification Trees

The main objective of classification trees is to classify the data according to different criteria. The dataset is continuously subdivided according to the characteristics of the data and each consequent subdivision depends on the value of the previous subdivision. The procedure is terminated when a class is finally predicted for the observations. The starting point is called the root node. The sequence starts with the entire learning set at the root node, and terminates in a terminal node which gives the final class label for the observations present at that node. The term CART which stands for Classification and Regression Trees was first introduced by Leo Breiman.

#### 3.2.4 Artificial Neural Networks

Artificial Neural Networks or ANNs are connectionist systems, based upon the connectionism theory of how the brain works in living organisms. The neural network is a method under which many machine learning algorithms work to process complex data. A neural network works in a manner similar to a biological neuron, where signals travel via an electrochemical process.

The brain learns by changing strengths of the connections between neurons or by adding or removing connections.

Similar to the biological system, an artificial neural network or ANN is based on a collection of connected units or nodes called artificial neurons, which work in a manner similar to the brain. Each connection, like the synapses in a biological brain, can transmit a signal from one artificial neuron to another. An artificial neuron that receives a signal can process it and then further transmit a signal to the other artificial neurons connected to it.

In ANN implementations, the signal at a connection between artificial neurons is a real number, and the output of each neuron is computed by some non-linear function of the sum of its inputs. Artificial neurons use a weight that adjusts during the process of learning. The weight may increase or decrease the strength of the signal at a connection. Artificial neurons may have a threshold such that the signal is only sent if the aggregate of the signal from all the neurons connected to a particular synapse crosses that threshold. Typically, artificial neurons are aggregated into layers. Signals travel from the first layer called the input layer, to the last layer called the output layer.

An ANN uses an activation function to check the state of the neurons, and using several different algorithms, it calculates the weights to be assigned to each independent variable. Some bias term is introduced at each level in order to generalize the result and the fitted model is obtained.

#### 3.2.5 Support Vector Machines

A Support Vector Machine (SVM) is a supervised learning technique which uses a separating hyperplane to classify the given data. It uses an algorithm to divide the classes for the given labelled training data according to an optimal hyperplane.

An SVM model represents the data points in space, and divides the points according to the different classes, using a suitable hyperplane. Then, new data points are classified accordingly and the predicted values are compared with the actual values to determine the accuracy of the model.

The support vectors in a support vector machine are the points which define the hyper plane. These points lie on the hyper plane. The classification rule for SVM depends on these support vectors.

#### 3.2.6 Ridge Regression

Ridge regression is a technique that is used to analyse data that suffers from multicollinearity. Multicollinearity is a phenomenon where some almost linear relationship may exist among the independent variables. It means that a certain predictor variable can be expressed in terms of the others, in a fairly accurate manner. The overall reliability of the predictive model may not change, but the individual coefficient estimates may show a lot of variation in response to small changes in the data. Hence, even though the model as a whole may predict the output variable, but it is difficult to check for the individual predictors.

For any data where multicollinearity occurs, the least squares estimates may be unbiased but the variances are large, so there may be a lot of deviation from the true value. A bias term is added to the regression estimates, so as to reduce the standard errors and obtain reliable estimates.

For Ridge Regression the variance inflation factor (VIF) is considered. The cut off value for VIF is 10, hence the value of R-squared will be 0.90. So, for the situations when the value of R-squared is greater or equal to 0.90, multicollinearity is present. So, under the process of ridge regression, a small value k is added to the diagonal elements of the correlation matrix. The bias and the variance covariance matrix are calculated.

There are different methods of determining k, such as from the ridge trace plot.The ridge trace plot is a plot of the ridge regression coefficients as a function of k. The value of k for which the regression coefficients are stable is selected. The smallest possible value of k is chosen, in order to minimize the bias. Other methods for the selection of k include those given by Hoerl, Kennard and Baldwin and the method given by Lawless and Wang.

#### 3.2.7 Least Absolute Shrinkage and Selection Operator

The least absolute shrinkage and selection operator (LASSO) is a regression analysis method that helps in variable selection in order to simplify the interpretation of the data. When the number of independent variables is reduced, the accuracy of the fitted model increases and it becomes easier to interpret the model.

The LASSO technique was initially introduced in the context of least squares. Using this technique, a subset of the independent variables is selected which are used in fitting the final model. This helps to reduce the complexity of the model and in turn, the data. The variables or covariates included in this subset are the ones which have a strong influence on the output variable.

The main difference between ridge regression and LASSO is the fact that while ridge regression can reduce overfitting and hence improve the prediction error, it does not make the data easier to interpret by reducing the dimensions of the set of independent covariates. LASSO sets some of the coefficients equal to zero, and thus performs both model selection as well as parameters estimation simultaneously. However, Ridge regression is easy to solve, but LASSO involves more complex computations.

## 4. RESULTS

### 4.1 Logit Model

The table for the misclassification rate is given as:

**Table 1.**

| **OBSERVED** | **PREDICTED** |  |  |
| --- | --- | --- | --- |
|  | **0(NOT**<br>**HAVING**<br>**DIABETES)** | **1(HAVING**<br>**DIABETES)** | **TOTAL** |
| **0(NOT**<br>**HAVING**<br>**DIABETES)** | 182 | 9 | 191 |
| **1(HAVING**<br>**DIABETES)** | 55 | 62 | 117 |
| **TOTAL** | 237 | 71 | 308 |

In this table, 182 patients who do not have diabetes are correctly classified as not having diabetes, 9 patients are misclassified as suffering from diabetes when they do not have diabetes. Similarly 55 patients are misclassified as not suffering from diabetes when they actually do suffer from diabetes, and 62 patients are such who are correctly classified as having diabetes when they do actually have diabetes.

The misclassification rate is obtained as **0.2077922** ≈ **21%.**

### 4.2 Probit Model

The table for the misclassification rate is obtained as

**Table 2.**
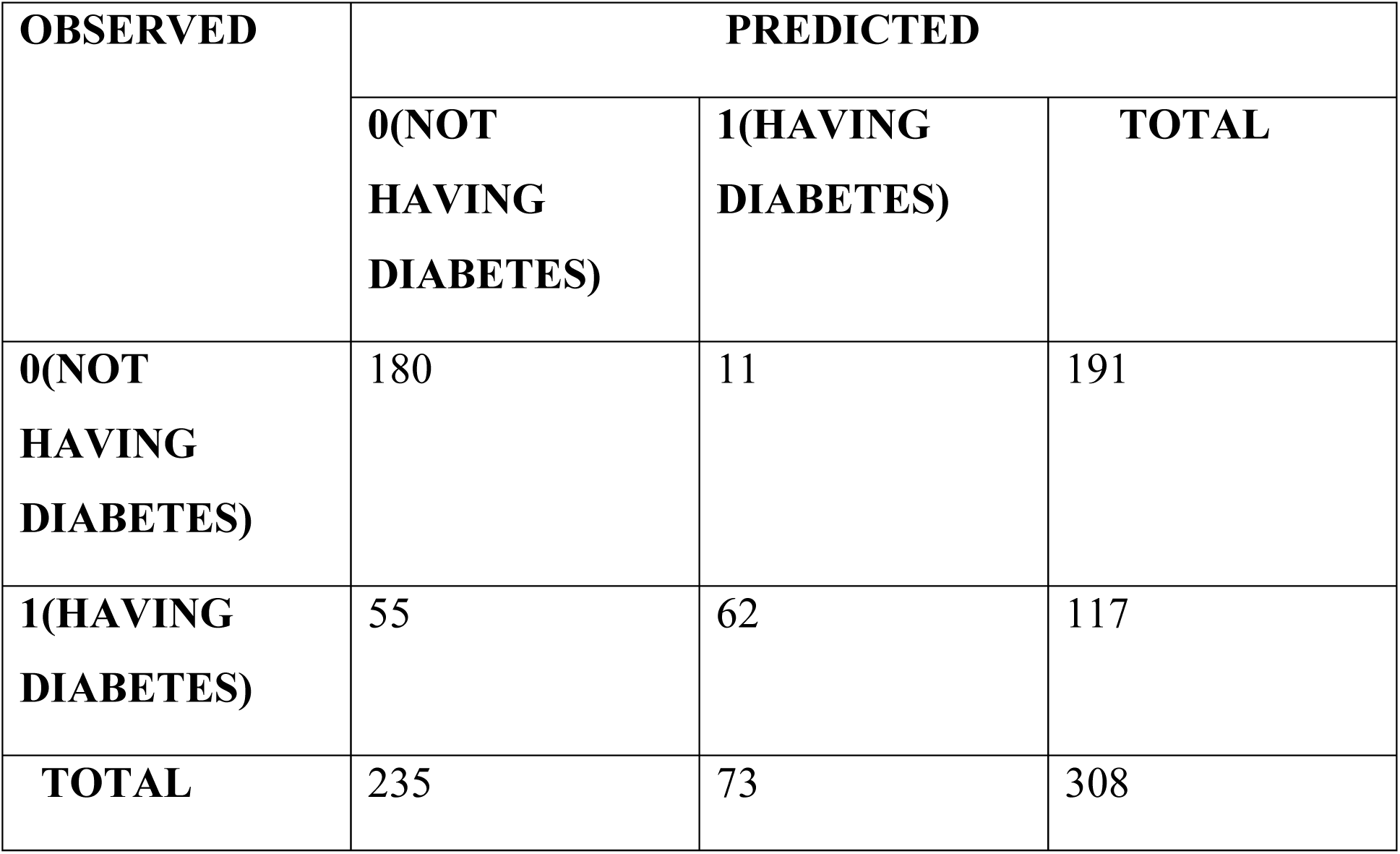

| **OBSERVED** | **PREDICTED** |  |  |
| --- | --- | --- | --- |
|  | **0(NOT**<br>**HAVING**<br>**DIABETES)** | **1(HAVING**<br>**DIABETES)** | **TOTAL** |
| **0(NOT**<br>**HAVING**<br>**DIABETES)** | 180 | 11 | 191 |
| **1(HAVING**<br>**DIABETES)** | 55 | 62 | 117 |
| **TOTAL** | 235 | 73 | 308 |

In this table, 180 patients who do not have diabetes are correctly classified as not having diabetes, 11 patients are misclassified as suffering from diabetes when they do not have diabetes which is more than the case for logit model. Similarly 55 patients are misclassified as not suffering from diabetes when they actually do suffer from diabetes, and 62 patients are such who are correctly classified as having diabetes when they do actually have diabetes.

The misclassification rate is obtained as **0.2142857** ≈ **21%.**

### 4.3 Classification Tree

Figure 1 shows the classification tree obtained for the training dataset.

**Fig. 1.**
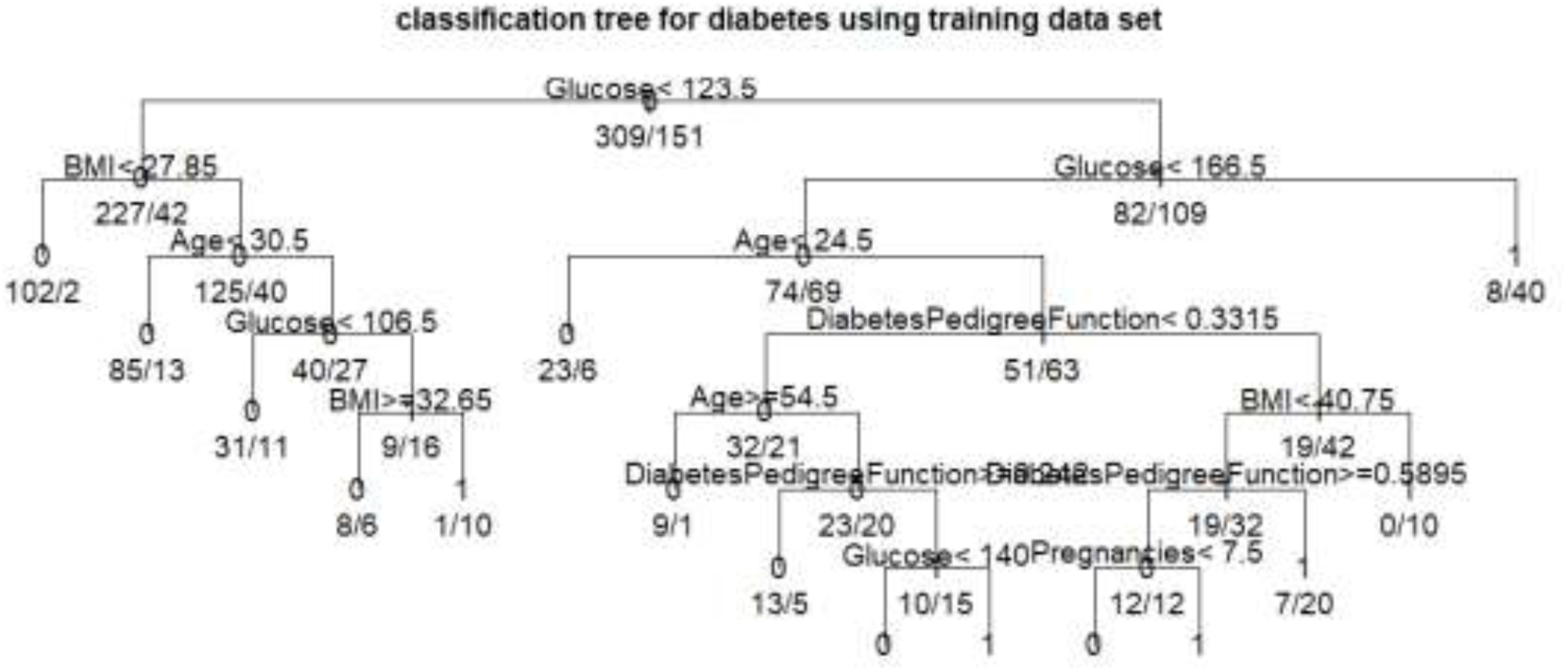

This classification tree has been constructed using the training dataset which contains 460 observations selected at random from the population.

Here the value of Glucose is initially considered as less than or greater than 123.5. For those 309 individuals for whom the value of Glucose is less than 123.5, their body mass index is further observed as to being less or greater than 27.85. For those 151 individuals for whom the value of Glucose is greater than 123.5, they are further classified by checking for a higher value of Glucose, that is, 166.5. Here 0 denoted that node at which the patients are classified as not suffering from diabetes, and 1 denotes the node at which the individuals are classified as suffering from diabetes. Hence, for those individuals whose Glucose level is higher than 166.5, they are classified as suffering from diabetes on the basis of high glucose content in the blood. For those patients for whom the value of glucose variable is less than 166.5, the next factor which is age is checked. By continuing this process, all the individuals in the training dataset are classified as suffering from diabetes or not on the basis of the different factors.

The table for the misclassification rate is obtained as

**Table 3.**

| **OBSERVED** | **PREDICTED** |  |  |
| --- | --- | --- | --- |
|  | **0(NOT**<br>**HAVING**<br>**DIABETES)** | **1(HAVING**<br>**DIABETES)** | **TOTAL** |
| **0(NOT**<br>**HAVING**<br>**DIABETES)** | 173 | 18 | 191 |
| **1(HAVING**<br>**DIABETES)** | 68 | 49 | 117 |
| **TOTAL** | 241 | 67 | 308 |

Here 173 patients who do not have diabetes are correctly classified as not having diabetes, whereas 18 patients are misclassified as suffering from diabetes when they do not have diabetes. Similarly 68 patients are misclassified as not suffering from diabetes when they actually do suffer from diabetes, and 49 patients are such who are correctly classified as having diabetes when they do actually have diabetes.

The misclassification rate is obtained as **0.2792208** ≈ **28%.**

### 4.4 Artificial Neural Network

The neural network that has been fitted for the training dataset is given in Figure 2.

**Fig. 2.**
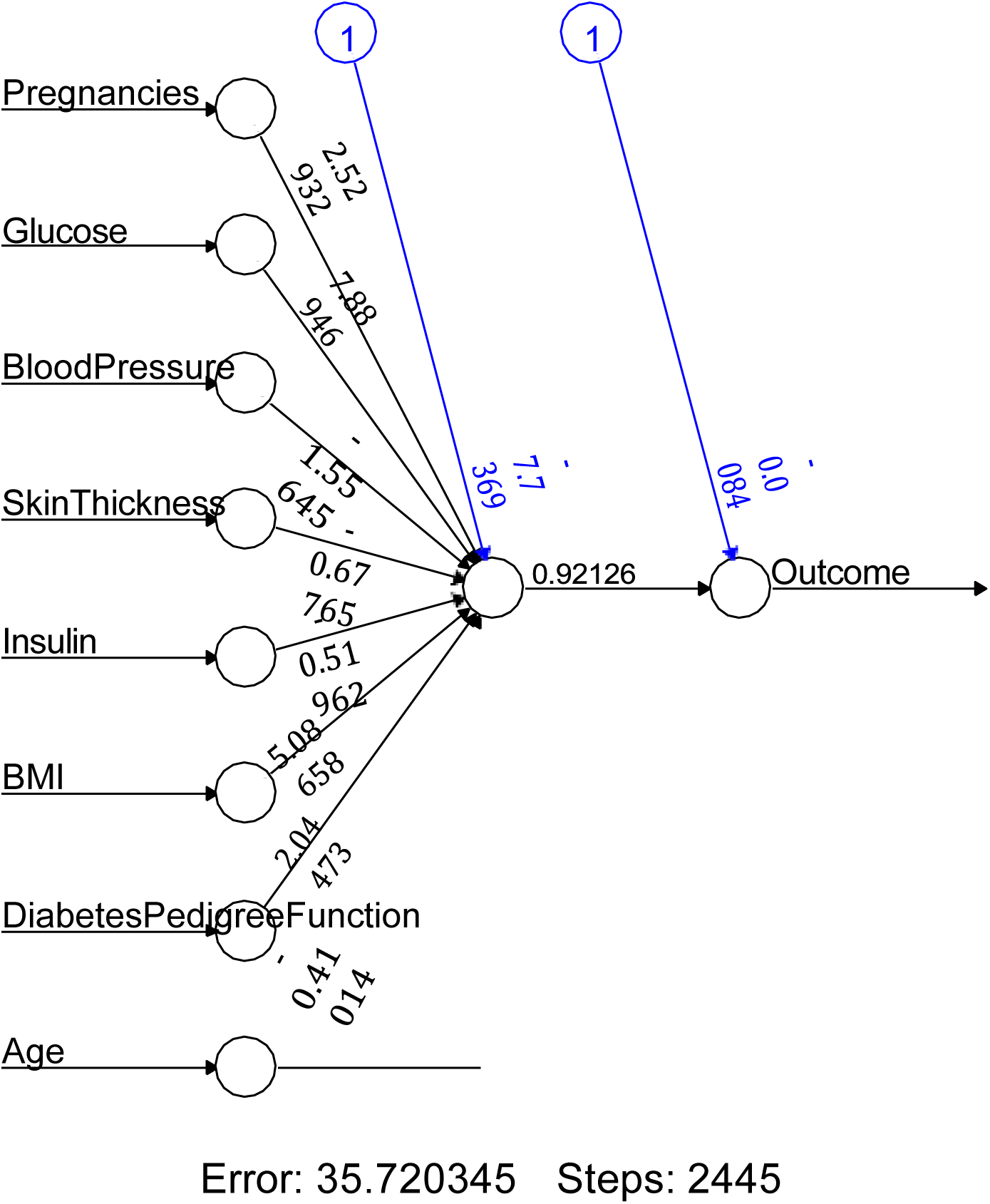

Figure 2 shows an artificial neural network with an input layer which consists of eight independent variables, a single hidden layer and an output layer. Here the weights have been recursively calculated using the back propagation algorithm, and a bias term has been introduced for the node in the hidden layer. This bias term enables the results to be generalized, instead of being limited to that particular instance of the data. The individual weights represent the strength of connections between the units. The activation function used here is the “logistic” activation function, in order to take into consideration any non linear properties in the network. Since the output is in the form of zero and one, that is, binary, therefore use of logistic regression is more appropriate. The reason for the use of back propagation algorithm is that it helps to calculate a gradient, which gives the direction of the greatest rate of increase of the function. It’s easier and more accurate to recursively calculate the weights, as compared to feedforward network where the data only moves in one direction, that is, forward.

The table for the misclassification rate for the ANN is obtained as

**Table 4.**

| **OBSERVED** | **PREDICTED** |  |  |
| --- | --- | --- | --- |
|  | **0(NOT**<br>**HAVING**<br>**DIABETES)** | **1(HAVING**<br>**DIABETES)** | **TOTAL** |
| **0(NOT**<br>**HAVING**<br>**DIABETES)** | 178 | 13 | 191 |
| **1(HAVING**<br>**DIABETES)** | 55 | 62 | 117 |
| **TOTAL** | 233 | 75 | 308 |

Here 178 patients who do not have diabetes are correctly classified as not having diabetes, whereas 13 patients are misclassified as suffering from diabetes when they do not have diabetes. Similarly 55 patients are misclassified as not suffering from diabetes when they actually do suffer from diabetes, and 62 patients are such who are correctly classified as having diabetes when they do actually have diabetes.

The misclassification rate is obtained as **0.2207792208** ≈ **22%.**

### 4.5 Support Vector Machine

The table for the misclassification rate for the SVM is obtained as

**Table 5.**

| **OBSERVED** | **PREDICTED** |  |  |
| --- | --- | --- | --- |
|  | **0(NOT**<br>**HAVING**<br>**DIABETES)** | **1(HAVING**<br>**DIABETES)** | **TOTAL** |
| **0(NOT**<br>**HAVING**<br>**DIABETES)** | 182 | 9 | 191 |
| **1(HAVING**<br>**DIABETES)** | 66 | 51 | 117 |
| **TOTAL** | 248 | 60 | 308 |

In the given table 182 patients who do not have diabetes are correctly classified as not having diabetes, whereas 9 patients are misclassified as suffering from diabetes when they do not have diabetes. Similarly 66 patients are misclassified as not suffering from diabetes when they actually do suffer from diabetes, and 51 patients are such who are correctly classified as having diabetes when they do actually have diabetes.

The misclassification rate is obtained as **0.2435064935**≈ **24%.**

### 4.6 Ridge Regression

The table for the misclassification rate is obtained as

**Table 6.**

| **OBSERVED** | **PREDICTED** |  |  |
| --- | --- | --- | --- |
|  | **0(NOT**<br>**HAVING**<br>**DIABETES)** | **1(HAVING**<br>**DIABETES)** | **TOTAL** |
| **0(NOT**<br>**HAVING**<br>**DIABETES)** | 185 | 6 | 191 |
| **1(HAVING**<br>**DIABETES)** | 70 | 47 | 117 |
| **TOTAL** | 255 | 53 | 308 |

In the given table 185 patients who do not have diabetes are correctly classified as not having diabetes, whereas 6 patients are misclassified as suffering from diabetes when they do not have diabetes. Similarly 70 patients are misclassified as not suffering from diabetes when they actually do suffer from diabetes, and 47 patients are such who are correctly classified as having diabetes when they do actually have diabetes.

The misclassification rate is obtained as **0.2467532468**≈ **25%.**

### 4.7 Least Absolute Shrinkage and Selection Operator

The table for the misclassification rate for LASSO is obtained as

**Table 7.**

| **OBSERVED** | **PREDICTED** |  |  |
| --- | --- | --- | --- |
|  | **0(NOT**<br>**HAVING**<br>**DIABETES)** | **1(HAVING**<br>**DIABETES)** | **TOTAL** |
| **0(NOT**<br>**HAVING**<br>**DIABETES)** | 186 | 5 | 191 |
| **1(HAVING**<br>**DIABETES)** | 72 | 45 | 117 |
| **TOTAL** | 258 | 50 | 308 |

Here 186 patients who do not have diabetes are correctly classified as not having diabetes, whereas 5 patients are misclassified as suffering from diabetes when they do not have diabetes. Similarly 72 patients are misclassified as not suffering from diabetes when they actually do suffer from diabetes, and 45 patients are such who are correctly classified as having diabetes when they do actually have diabetes.

The misclassification rate is obtained as **0.25** ≈ **25%.**

## 5. CONCLUSION

Gestational diabetes has a detrimental effect not only the future health of the mother, but also on the health of her child, who may become more prone to suffering from diabetes in later life. For these reasons, it is important to develop a prediction model with maximum accuracy in order to predict on the basis of observed factors, the chances of an individual developing diabetes.

A model with maximum accuracy will be that model for which the classification of patients as suffering from diabetes or not suffering from diabetes is the closest to the actual values. Hence, that method of classification will be considered the best which has the least misclassification rate.

In this study, 7 different methods have been considered, for which the misclassification rates are obtained as follows:

**Table 8.**

| **Method** | **Misclassification rate in %** |
| --- | --- |
| Logit Model | $0.2077922 \approx 21\%$. |
| Probit Model | $0.2142857 \approx 21\%$. |
| Classification Tree | $0.2792208 \approx 28\%$. |
| Artificial Neural Network | $0.2207792208 \approx 22\%$. |
| Support Vector Machine | $0.2435064935 \approx 24\%$. |
| Ridge Regression | $0.2467532468 \approx 25\%$. |
| LASSO | $0.25 \approx 25\%$. |

It can be seen that the logit model has the minimum misclassification rate out of all the seven methods applied to the data. Hence, it can be deduced that the logit model may be considered as the most suitable model to be used for this study.

Further, it is seen that LASSO assigns importance to three independent variables, that is, Pregnancies, which denotes the number of pregnancies for the patients, Glucose, which denotes the amount of glucose in the body of the patients, and BMI which shows the body mass index of the patient, that is, whether the patient is overweight or underweight. Hence it can be said that these three variables are the most important out of the eight independent variables in determining whether or not the patients suffer from diabetes, that is, a subset of three variables has been obtained from the set of eight independent variables. This is a major benefit of using LASSO.

## 6. DISCUSSION

It has been observed in the study that the classification tree technique has the worst performance, since the misclassification rate is nearly 28%, whereas the logit model has the best performance with a misclassification rate of approximately 21%. Since the logit and the probit model are both suitable for the classification of binary data, and differ in the use of the link function, where the logit model uses a logit link function and the probit model uses an inverse normal link function, therefore there is very little difference in the misclassification rates obtained by using the logit and the probit models. After the logit and the probit models, the artificial neural network is seen to be the next most suitable method for classifying the given dataset, followed by the support vector machine, the ridge regression technique, and then LASSO method. The difference between ridge and LASSO's misclassification rates may be accounted for because of the penalties which are imposed on the data under these models. The ridge regression technique applies the ***l_2_*** penalty, whereas the LASSO technique applies the ***l_1_*** penalty.

